# A stochastic ratchet during genome streamlining can commit insect endosymbionts to a parasitic or mutualistic fate

**DOI:** 10.1101/2025.04.10.648247

**Authors:** Pranas Grigaitis, Bas Teusink, Adria C. LeBoeuf

**Affiliations:** Systems Biology Lab, Amsterdam Institute for Life and Environment (A-LIFE) & Amsterdam Institute for Molecular Life Sciences (AIMMS), Vrije Universiteit Amsterdam, Amsterdam, the Netherlands; Department of Zoology, University of Cambridge, United Kingdom

**Keywords:** endosymbiotic bacteria, symbiosis, metabolic capacity, genome streamlining, cooperation, genome-scale metabolic modeling

## Abstract

Many arthropods, including insects, harbor endosymbiotic bacteria. As bacteria transitioned to live inside host cells, free-living bacterial ancestors of endosymbionts reduced their genomes in a process known as streamlining. Contemporary endosymbioses represent at least two distinct life strategies: endosymbionts either become unilaterally dependent on host-derived resources (uni-obligate) or enter bidirectionally obligate mutualistic relationships (bi-obligate endosymbiosis). It is an open question whether these paths taken by free-living bacteria can be sequential, or alternatively, if commitment takes place early, making these strategies mutually exclusive. In this study, we quantified metabolic capacity of different endosymbionts to contextualize genome streamlining in terms of organismal function. We show that uni-obligate endosymbionts of insects have lost substantial parts of their metabolic capacity that are present both in free-living bacteria and bi-obligate endosymbionts. On the contrary, genomes of uni-obligate endosymbionts contain more genes which can be attributed to scavenging host resources. In sum, we challenge the existing notion that bi-obligate endosymbionts have emerged from uni-obligate ones, and rather, propose a stochastic ratchet model: in early stages of endosymbiosis, bacteria get locked into either a more mutualistic or more parasitic relationship with the host depending on the first losses during streamlining.

## Introduction

Most life engages in relationships with other species that fall along a spectrum from bidirectionally beneficial (mutualistic) to one-sidedly positive at the detriment of the other partner (parasitic) [1]. Many of these interactions are formed between and/or with microbes, frequently in niches created by multicellular eukaryotes, for example, in animal gastrointestinal- and reproductive systems, or in association with plant root nodules. An extreme case of such an association between microbes and multicellular eukaryotes is endosymbiosis, or intracellular colonization of host cells by microbes.

From the microbial perspective, engulfment of free-living bacteria in a nutrient-rich host cell’s cytosol creates situation where (i) effective population size is small (increased genetic drift), and (ii) nutrition-related selection pressures are alleviated. As a consequence, entering the endosymbiotic lifestyle drastically accelerates the evolution of their genomes [2,3] and many endosymbiont genomes are reduced via genome streamlining [4]. At the infancy of the endosymbiotic relationship, bacterial genomes typically lose parts of the genome that are redundant with the host’s genome. Such a Muller’s rachet [5] effectively prevents their reversal to the free-living lifestyle [6], and these bacteria develop single-sided dependency on the host (become uni-obligate).

One animal lineage that has frequently become associated with different microbes are the insects [7]. In endosymbiosis, like in any collaboration, there is conflict (misalignment of fitness goals between partners). Such conflict can be managed through the evolution of conflict meditation strategies in both partners [8]. For example, some contemporary uni-obligate insect endosymbionts promote their own transmission by altering the sex ratio of host brood [9] while hosts evolve to counter these effects [8,10]. Successful cooperative endosymbionts can offer massive benefits to the host, e.g. enabling hosts to colonize novel ecological niches via supplementation of missing nutrients like essential amino acids [11,12] or B vitamins [13]. In such cases, both parties sometimes co-evolve into mutually obligate (bi-obligate) relationship, culminating in a major evolutionary transition in individuality (METI) [14].

Compared to the oldest known endosymbionts (ancestors of mitochondria and chloroplasts), insect endosymbiotic relationships are recent (emerged <300 Mya [6]) and are still undergoing development [15]. The origins of endosymbioses are under ongoing debate; it has been suggested that parasitic free-living bacteria evolved to reduce their virulence and enabling co-habitation with a host [16] and host-mediated transmission [17]. How do microbes commit to either uni- or bi-obligate endosymbiosis? Currently, uni-obligate endosymbiosis is treated as an intermediate evolutionary state in a linear path for free-living bacteria to become bi-obligate [18,19]. However, theory in the domain of METIs would suggest that once conflict mediation mechanisms take root to manage significant conflicts of interest (as eluded for uni-obligate endosymbionts above), major evolutionary transitions are inhibited [20]. We hypothesize that genome streamlining towards uni- and bi-obligacy could be two mutually exclusive evolutionary paths rather than a sequential process, with stochastic, irreversible decision-making taking place early on, which we term a stochastic ratchet.

Here we challenge the linear commitment model by mapping the genome content of different endosymbiotic bacteria to their metabolic capacity. To test the stochastic ratchet hypothesis, we quantified the metabolic capacity of over 500 uni- and bi-obligate insect endosymbionts. We reconstructed genome-scale metabolic models (GEMs) [21] based on genome sequences and found that, in general, the metabolic capacity of bi-obligate endosymbionts was higher than uni-obligate endosymbionts despite bi-obligate genomes being more strongly reduced in size. Our work highlights different strategies endosymbiotic bacteria followed that are mutually incompatible to form a chain of successive increase of commitment from lifestyle of free-living bacteria to obligate endosymbiosis.

## Results

### Endosymbiont genomes have undergone various degrees of reduction

We have collected 542 genomes of endosymbiotic bacteria from five genera: Candidatus *Blochmanniella* (hereafter *Blochmanniella*), *Buchnera, Rickettsia, Spiroplasma*, and *Wolbachia* (Methods, Table S1, Figure S1). These microbial genera can be classified into bi-obligate mutualistic (*Blochmanniella, Buchnera*), and uni-obligate endosymbionts (*Rickettsia, Spiroplasma*, and *Wolbachia)*; in the latter group, the endosymbiont has a context-dependent impact on the host, ranging from mutualism to parasitism. We chose not to include genera which are culturable on axenic media [22] and/or have genome sizes comparable to free-living bacteria (Figure S1) like *Sodalis, Serratia*, or *Carsonella*. We first compared the genome sizes and content of open reading frames (ORFs) across the genera.

Around half of the genomes in our cohort were *Wolbachia*, and all sixty-eight *Buchnera* samples were isolates of a single species, *B. aphidicola*. For all genera, the number of open reading frames (ORFs) showed linear increase with increasing genome size (R^2^ = 0.893, Figure 1a, Figure S1). The largest genome, *Rickettsia* endosymbiont of *Oedothorax gibbosus* (GCF_936269705.1) was 2.6 Mbp (2915 ORFs). Consistent with previous observations [23], the genomes of bi-obligate mutualist endosymbionts were smaller than those of uni-obligate endosymbionts (Figure 1a; U-test, p<0.01).

**Figure 1.**
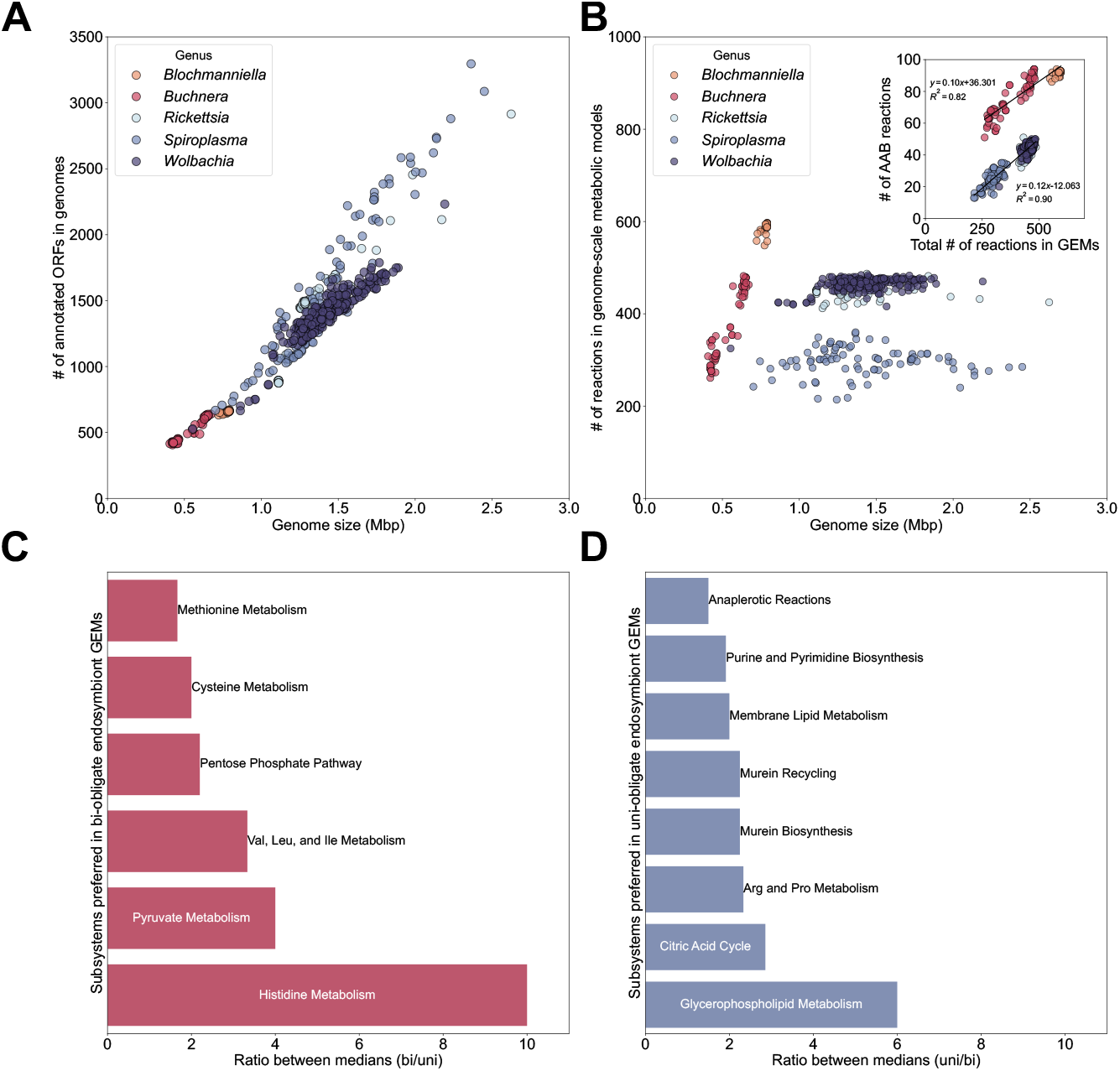
Endosymbiont genome size does not correlate with the number of metabolic genes. **a**. The number of open reading frames (ORFs) as a function of genome size in different genera. **b**. The number of reactions in the resulting draft GEMs. Inset of (b): Number of metabolic reactions related to amino acid biosynthesis in draft GEMs (see Methods). **c-d**. Preference scores (median ratios) for presence of reactions in GEM subsystems (see Methods), with values in (c) representing preference for bi-obligate *Blochmanniella* and *Buchnera*, and in (d) for uni-obligate *Rickettsia, Spiroplasma*, and *Wolbachia* endosymbiont GEMs. In (c) and (d), only subsystems with median ratio >1.5-fold and significantly different content in uni- and bi-obligate endosymbionts (U-test, p<0.05) are shown.

To put the genomic metrics into the context, we did comparisons against two representative bacterial genera: a pathogenic intracellular bacterium *Mycoplasma* and a free-living bacterium *Escherichia coli*. All endosymbiont genomes were considerably smaller than an *E. coli* MG1655 (4.6 Mbp, 4639 ORFs, genome assembly GCF_000005845.2) (Figure S1). Bi-obligate endosymbionts had smaller genomes than *Mycoplasma*, and the opposite was true for uni-obligate endosymbionts (Figure S1; U-test, p<0.01 for both comparisons). This places *Mycoplasma*, a parasitic bacterium with a broad spectrum of hosts (known to infect multiple animals and plants [24]) apart from the endosymbiotic bacteria found in associations with insects. Combined, our observations confirm that genomes of endosymbiotic bacteria have undergone various degrees of genome streamlining, likely driven by their evolutionary and ecological history.

### Endosymbiont genome size does not correlate with the number of metabolic genes

Since bi-obligate endosymbionts maintain smaller genomes with fewer ORFs than the uni-obligate endosymbionts, we asked whether genomes of bi-obligate endosymbionts also harbor fewer metabolic genes and have reduced metabolic capacity. We reconstructed genome-scale metabolic models (GEMs) for all endosymbiont genomes using the GEM of *Escherichia coli* as the template. This method is used to to reconstruct a metabolic network of an organism-of-interest by matching ORFs with known metabolic function from the template genome with sequences in the target genome sequence. We excluded all the transport reactions (and transporter genes, Figure S2) from the analysis of draft models since many metabolite-transporter pairs are unknown even in a well-studied organism like *E. coli* [25]. Moreover, host-endosymbiont metabolite transport is poorly understood, and we chose to avoid bias in evaluating metrics stemming from ambiguous annotation of transporters.

To our surprise, we observed that GEMs of bi-obligate endosymbionts harbor more metabolic reactions than uni-obligates despite their smaller genomes (Figure 1b), as did the models including transport reactions matched with the template model (Figure S3). We found that *Blochmanniella* GEMs contained the most metabolic reactions of the tested genera. *Buchnera*-uni-obligate endosymbiont GEMs contained similar number of reactions (*Buchnera* vs. *Wolbachia* or *Rickettsia*; in both cases, U-test p>0.05), however, *Buchnera* was the only genus where the number of reactions linearly increased with genome size (R^2^ of 0.92). the GEMs of bi-obligate endosymbionts *Blochmanniella* and *Buchnera* had more amino acid biosynthesis reactions compared to uni-obligates (Figure 1b, inset). The number highly correlated with number of reactions in the GEM of uni-obligates (R^2^ of 0.89 and 0.82 for uni- and bi-obligates, respectively) but not with genome size (R^2^ of 0.007 vs 0.80 for uni- and bi-obligates, respectively).

We compared the number of reactions in pathways (subsystems) in respective GEMs to infer differential representation of metabolic pathways in uni-vs. bi-obligate endosymbionts (Figure 1c, 1d). We computed ratios of median number of reactions between the groups and found that reactions for amino acid biosynthesis are more preferred in the GEMs of bi-obligate endosymbionts, while reactions for cell envelope biosynthesis and catabolic processes were preferred in uni-obligate endosymbionts. Overall, our results hint into different functions of bi-obligate vs. uni-obligate metabolic networks (bottom-up biosynthesis of cell components vs. assembly of pre-made components into new cells).

### Endosymbiont genomes and metabolisms are highly variable

We also wanted to quantify the intra-genus genome variability [26] in our cohort of genomes. For this, we clustered the ORFs of individual genera into groups of orthologous genes, or orthogroups (Figure S5). Consistent with previous observations, we found that many orthogroups were present in only a few genomes within the genus. We next asked whether the variation in content of endosymbiont genomes was reflected in respective GEMs.

The GEMs of same genus contained similar reaction sets, as suggested by principal component analysis (Figure S3). Clustering was evident in the first two principal components that together explain almost 60% of total variance. However, only seven metabolic reactions were present in all 542 GEMs reconstructed. Even reactions critical for producing new cell biomass, e.g. amino acyl-tRNA loading, were not consistently present in all GEMs (Figure S4). Thus, to compare the metabolic networks in an analogous fashion to the orthogroups, we next computed pan-GEMs: genus-level unions of metabolic reactions. This entailed collecting all reactions present in at least one GEM within the genus (Figure S5). Of the 1004 metabolic reactions present in at least one individual GEM, only 152 (15%) of them were shared by all five genus-level pan-GEMs (Figure S6); most of these reactions metabolize host-derived biomass precursors and represent the capacity of endosymbionts to assimilate host-derived precursors.

### Endosymbiont metabolic networks contain gaps to be filled by the host

GEMs, or metabolic network reconstructions, are databases of all metabolic reactions that an organism can perform given the enzymes encoded in their genome. However, presence of individual reactions does not directly translate in metabolic capacity, which is determined by functional pathways. We compiled a set of more than 40 biomass precursors (amino acids, nucleotides, and intermediates of central carbon metabolism, Table S3) to test whether the GEMs could produce biomass precursors from nutrients supplied by the host (Figure 2, Figure S7).

**Figure 2.**
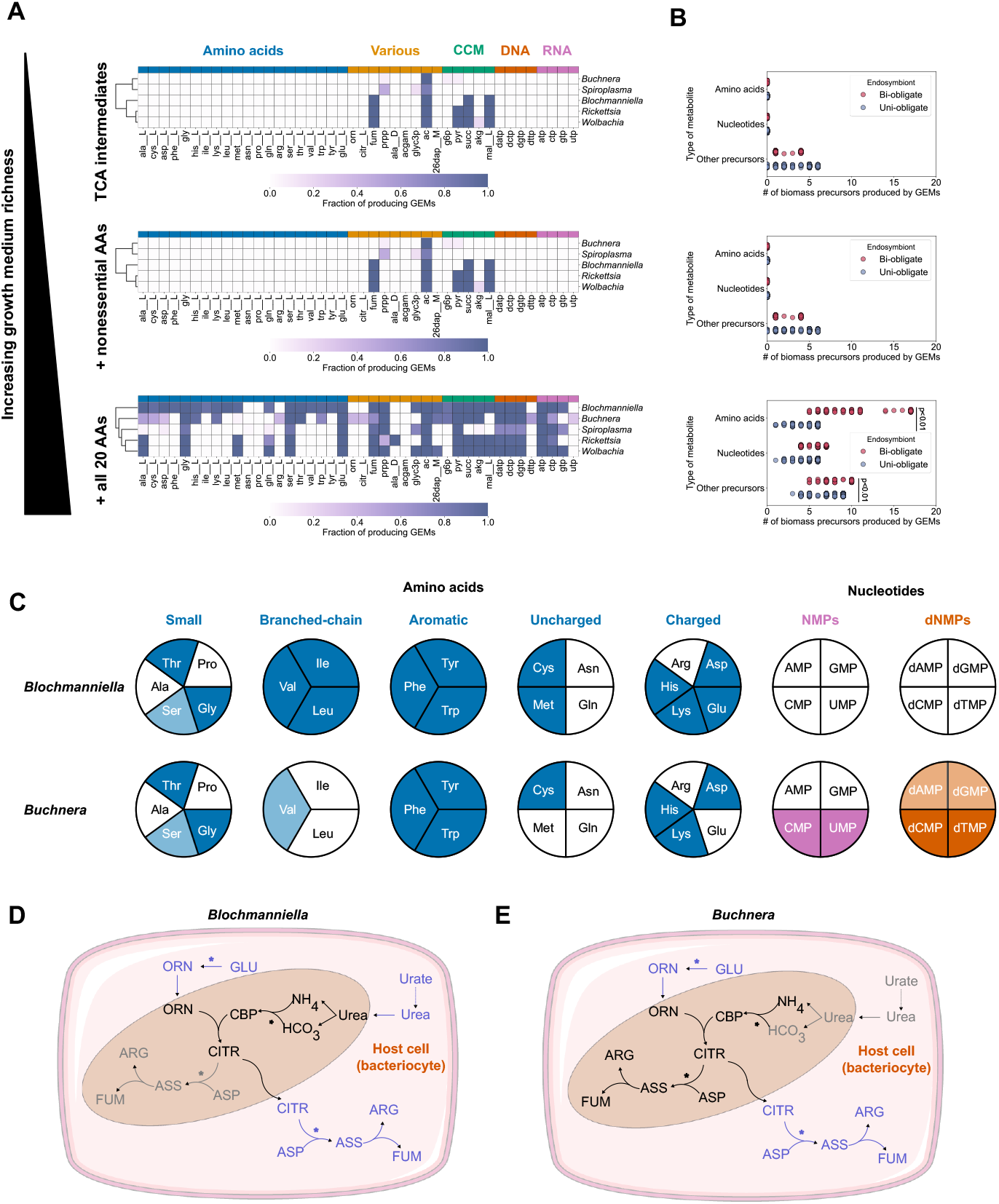
Amino acids are primary metabolic goods produced by bi-obligate endosymbiont metabolic networks. **a-b**. Production analysis for endosymbiont GEMs. a. The intensity of the color represents the fraction of GEMs per genus, capable of producing a metabolite (see Methods). CCM, central carbon metabolism. **b**. Number of produced metabolic precursors (amino acids, nucleotides, or other intermediates) in each GEM. In (b), GEMs are grouped per obligacy status (uni-/bi-obligate), and “other intermediates” category represents the sum of “Various” and “CCM” metabolites in panel (a). **c-e**. Biosynthetic capacity of representative, manually curated *Blochmanniella* and *Buchnera* GEMs with organic acids and inorganic nitrogen as carbon- and nitrogen sources, respectively. **c**. Manual analysis of metabolic capacity in GEMs. Dark-shaded blocks represent complete biosynthesis pathways, and lighter shade represents partial biosynthesis. **d-e**. Differences in urea assimilation and arginine production pathways in *Blochmannia floridanus* (d) and *Buchnera aphidicola* (e). Missing conversions in the endosymbiont GEMs are marked in gray. Blue metabolites and arrows represent processes happening in the host cell, asterisks denote reactions hydrolyzing ATP. Abbreviations in (d) and (e): ORN, ornithine; GLU, L-glutamate; CBP, carbamoyl phosphate; CITR, L-citrulline; ASP, L-aspartate; ASS, arginino-succinate; FUM, fumarate, ARG, L-arginine.

We have simulated three scenarios representing different access to host-derived nutrients (Figure 2). Prior to simulation, we added all metabolic reactions to the GEMs from the template model that were not associated to any enzyme (see Methods), i.e. reactions for which no enzyme is known or needed. Moreover, due to the ambiguous nature of metabolite transport between host and endosymbiont, we assumed all metabolites in the GEM can be transported from and to the external (host) environment. The richness of the medium, from poor media (containing no organic nitrogen sources) to supplementation with all 20 amino acids (rich media) represents different levels of host cooperation in the production of biomass precursors in the endosymbiont GEMs. In all cases, the models were allowed to take up inorganic ions (ammonium, phosphate, sulfate) and oxygen; only carbon- and organic nitrogen sources differed among scenarios (Figure 2a).

Few metabolic intermediates could be produced in all endosymbiont GEMs on poor media (Figure 2b). The production capacity differed only for the nutrient-rich condition (all amino acids available): amino acid-and nucleotide biosynthesis capacity differs between uni- and bi-obligate endosymbionts (Figure 2b; U-test, p<0.01 for amino acids and nucleotides, p=0.13 for other intermediates). Most GEMs required supply from the host for cell wall biosynthetic precursors N-acetyl-D-glucosamine and D-alanine, and pyrimidine nucleotides. The most distinct difference across genera was that *Blochmanniella* and *Buchnera* GEMs could produce most amino acids (except, e.g., L-asparagine) (Figure 2b); on the contrary, most of the GEMs of uni-obligate endosymbionts could not produce any amino acids except for glycine and L-serine. In sum, these results suggest that the GEMs of uni-obligate endosymbionts cannot produce most biomass precursors, consistent with our previous observation that these GEMs contain fewer metabolic reactions than most GEMs of bi-obligate endosymbionts.

### The host engages in mutually beneficial metabolic division of labor with bi-obligate endosymbionts

Our previous analysis has suggested that host metabolism must complement the reduced metabolic networks of endosymbionts. It is well established that *Blochmanniella* and *Buchnera* supply amino acids to their hosts [11,27]. Yet the same functional outcome – non-trivially – can be achieved via differential division of labor between host- and endosymbiont metabolic networks. Based on the results of our production analysis (Figure 2b), we investigated host-endosymbiont complementarity more deeply by manually curating *Blochmanniella floridana* (*Bflo*) and *Buchnera aphidicola* (*Baph*) GEMs (Figure 2c).

The most striking differences between metabolisms of *Bflo* and *Baph* are differential capacity of assimilating nitrogen and producing nitrogenous compounds (Figure 2d, 2e). The manual curation resolved some artifacts in the automatically reconstructed GEMs, e.g., neither *Bflo* nor *Baph* produces L-serine (missing final step, phosphoserine phosphatase), or L-alanine from pyruvate (absent transaminase reaction) (Figure 2c). The *Bflo* metabolic network converts host-inaccessible N-sources (uric acid and urea) into accessible N (ammonium) using the enzyme urease. However, the *Bflo* urea cycle misses two final steps, and the cycle ends at L-citrulline. On the contrary, the *Baph* metabolic network contains a complete L-arginine synthesis pathway but not the urease reaction. *Bflo* could not produce nucleotides without host input, standing in contrast to the complete pyrimidine biosynthesis pathways in *Baph*. Overall, here we show that despite distinct modes of metabolic division of labor between *Bflo* or *Baph* and their hosts, their interactions with the host are reciprocally beneficial due to enrichment of host metabolism with essential amino acids missing in their N-poor diets.

### Genomes of uni-obligate endosymbionts retain genes to benefit from their hosts

The low metabolic gene content in genomes of uni-obligate endosymbionts and low metabolic capacity of their respective GEMs (Figure 1b) prompted us to look at what the functions of the “remaining” genes in the genome might be. For this, we inferred Gene Ontology (GO) terms for all 542 genomes, and searched for GO terms distinctively populated across genera. Many GO terms were represented only by very small number (<5) of genes in most of the genomes, and, to infer both biologically and statistically meaningful comparisons, we looked at GO terms with statistically significant differences between genomes of uni- and bi-obligate endosymbionts (U-test, p<0.01), and if more than 30 genes were assigned to the GO term in at least one genome (Figure 3).

**Figure 3.**
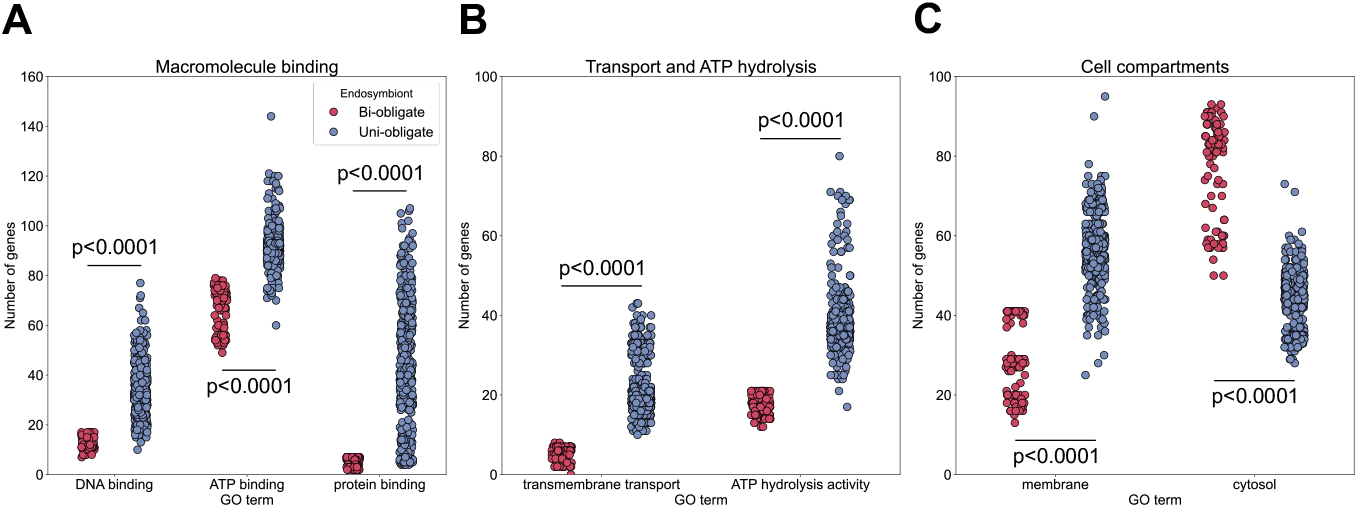
Uni-obligate endosymbiont genomes harbor more genes facilitating endosymbiont-host interactions and assimilation of host-derived nutrients. Distribution of genes in endosymbiont genomes based on their gene ontology (GO) terms, related to macromolecule binding (a), cell function and transport processes (b), and compartmentalization of proteins (c). All shown distributions of number of genes assigned to GO terms per genome were significantly different among individual genera (Kruskal-Wallis ANOVA, p<0.0001), and comparing groups of genera (uni-obligate vs. bi-obligate; U-test, p<0.0001).

We observed that genomes of *Rickettsia, Spiroplasma*, and *Wolbachia* had more genes annotated as “macromolecule binding” (Figure 3a). Moreover, the higher number of genes annotated as “transmembrane transport” and “ATP hydrolysis activity” in uni-obligate endosymbionts (Figure 3b) points to a higher capacity to transport and use host-derived precursors. Finally, the localization of gene products differentiated bi-obligate and uni-obligate endosymbionts: number of genes explicitly annotated as cytosolic (Figure 3c) was higher in *Blochmanniella* and *Buchnera* relative to the other genera, while we observed the opposite for genes associated with membrane localization. Some of these differences collapsed when the number of genes was normalized by the total number of genes in genomes (Figure S8), however, an absolute number of genes, and not fraction of total genome content, translates into presence of function. All in all, the genomes of *Rickettsia, Spiroplasma*, and *Wolbachia* seem to contain more genes that reflect key functions for a parasitic lifestyle.

## Discussion

Endosymbiotic bacteria colonize many arthropods, with multiple consequences of co-habitation for host physiology and ecology [14,16]. To deepen our understanding of endosymbiosis in terms of the metabolic interactions with the host, we reconstructed genome-scale metabolic models (GEMs) for 542 insect endosymbionts. Here we built on previous analyses of *Buchnera* [28], *Sodalis* [29], *Wolbachia* [30], and *Rickettsia* [31] GEMs at larger scale allowing to compare metabolic capacity across multiple genera of endosymbiotic bacteria. We found that the genomes of bi-obligate endosymbionts (genera *Blochmanniella* and *Buchnera*) contained more metabolic genes and functional pathways despite bearing more reduced (<1Mbp) genomes compared to their uni-obligate counterparts.

The overall host-endosymbiont economy always incurs host investments and only occasionally, metabolic returns from the endosymbiont [32]. In both cases, our results suggest that different processes lead endosymbionts to become obligate partners vs. obligate parasites. We observed a direct link between number of individual metabolic reactions in uni-obligate endosymbiont GEMs (especially of *Spiroplasma*, Figure 1b) and functional biosynthetic pathways (Figure 2a). The inferior metabolic capacity of uni-obligate endosymbionts is complemented by more genes that can be attributed to scavenging host resources (Figure 3, [33]). This way, we have established a firm, function-based distinction between different genome streamlining strategies that endosymbiotic bacteria have followed, one where metabolic capacity is prioritized for maintenance in the genome, and another where metabolic genes are consistently and stochastically lost (e.g., shikimate pathway genes [34]).

This is at odds with current understanding that bi-obligate endosymbiosis emerges out of uni-obligate endosymbiosis [33]. Once a uni-obligate endosymbiont has lost its metabolic capacity, it no longer has much to offer the host to create a mutually beneficial arrangement. Transitions to bi-obligate endosymbiosis are considered major evolutionary transitions in individuality [35,36]. Such transitions require a shift from individual priorities to collective priorities as the parliaments of genes of two organisms come together in what is termed an egalitarian major evolutionary transition [37]. In fraternal major evolutionary transitions (e.g., the transition to multicellularity), it has been suggested that when individual interests are sufficiently retained, resulting in both significant conflict and conflict mediation, this prevents transition to a new more complex organismal level [20]. In our system of endosymbiosis, once an endosymbiont has begun association with a host on a uni-obligate path, both a lack of host-relevant skills and a build-up of conflict and mediation strategies are likely to inhibit the transition to bi-obligacy.

Together, the stochastic loss of metabolic capacities that must come about under the pressure to streamline coupled with the gradual evolution of metabolic division of labor between host and endosymbiont provide the perfect opportunity for an evolutionary ratchet [38]. As the two partners specialize [39], certain genes are no longer in use and are lost stochastically (‘use it or lose it’), locking the endosymbiont into its collaboration with the host. As this allows hosts access to otherwise inhospitable diets [7], and consequently new niche spaces, they become reliant on the endosymbionts [40], ratcheting further toward a major evolutionary transition. Our work provides evidence for both this metabolic division of labor and for the stochastic loss of metabolic genes in egalitarian major evolutionary transitions. This model would also be consistent with bi-obligate endosymbiosis beginning as ectosymbiosis [41,42], where the two organisms might begin with a facultative mutualistic relationship before one is engulfed by the other.

An alternative explanation, a series of horizontal gene transfer (HGT) events that allow the genome of a uni-obligate endosymbiont to convert to the genome of a bi-obligate endosymbiont, is not supported by parametric metrics: sequences of bi-obligate-specific genes have comparable GC content to the content of aggregated coding sequences in respective genomes (Figure S8). Moreover, the hypothetical HGT events would involve massive rearrangements of the genome (replacing non-metabolic genes with metabolic ones, and reducing the resulting genome size several-fold, Figure 1).

Many essential parts of the physiology of endosymbiotic bacteria, such as synthesis and function of the cell wall [43,44], assembly of macromolecular complexes (nucleic acid polymerases and ribosomes) [45], and additional host benefits (e.g. apoptosis resistance [46]) remain an active object of study. Next, an endosymbiont can incur condition-dependent ecological effects: here we considered *Wolbachia* as a group of metabolic parasites but recent studies assign a spectrum of behaviors to different *Wolbachia* lineages [47–49]. The metabolic capacity of insects hosting bi-obligate endosymbionts is also an untapped asset, and diversity of combined host-endosymbiont metabolism is worth further exploring – with application(s) in mind [50,51]. To map genome content to metabolic function, here we applied canonical, primary protein structure-driven methods [52]. Degeneration of endosymbiont genomes, however, leads to many metabolic enzymes acquiring moonlighting functions [23,53] – to be addressed in future studies exploiting GEMs [23,53]. All in all, we believe that our study will pave the way for a more thorough follow-up investigation of metabolic cooperation, or division of labor, between insect endosymbionts and their hosts.

## Methods

### Collection of genomic data

We have collected the translated sequences of ORFs in genome sequences for genera of interest (*Blochmanniella, Buchnera, Wolbachia, Spiroplasma, Rickettsia*, and *Sodalis*) from the NCBI Datasets repository [54]. For reconstructing the draft GEMs, we downloaded all the genomes from NCBI Genome of these 6 genera satisfying the following criteria: (i) complete genomes; (ii) not atypical and metagenome-assembled genomes; (iii) genomes annotated by NCBI RefSeq. The search yielded 548 genome assemblies as of 1^st^ July, 2024 (Table S1). Functional annotation, including Gene Ontology term assignment, of detected ORFs was done with InterProScan 5.69-101.0 [55].

### Modification of Escherichia *coli* GEM iML1515

*Escherichia coli* GEM iML1515 [56] underwent minor modifications before its use as the template model for the endosymbiont GEMs. In the original GEM, only one amino acyl-tRNA loading reaction (for glutamate) was defined. We created reactions (and respective metabolites) for the remaining 19 proteinogenic amino acids. For these reactions, we assigned gene-protein-reaction associations from on the reference proteome of the *E. coli* strain MG1655, downloaded from UniProt [57].

### Automatic reconstruction of endosymbiont GEMs

The draft GEMs were generated using the template-based reconstruction routine, implemented in the RAVEN Toolbox 2.9.2 [52] under MATLAB 2023b environment [58]. The modified GEM iML1515-tRNA was used as the template model. The parameters for GEM reconstruction out of bi-directional BLAST results were set to their defaults except for minimal sequence alignment length (100 amino acids), protein BLAST e-value (10^−10^), and the sequence identity threshold (25%).

### Comparison of draft GEMs

First, all transport reactions were removed from comparison due to ambiguous representation in the template model and functionality in endosymbiotic bacteria. Then the union of all reactions in the draft models was taken and a binary matrix of presence (1) or absence (0) of a reaction was constructed.

Preference scores for subsystems of GEMs were determined by comparing the representing reaction counts in specific subsystem for bi-obligate vs. uni-obligate endosymbionts using the U-test. The pan-models of genera were created by taking the union of all metabolic reactions present in at least one GEM of that genus. Principal component analysis on the binary matrix computation was performed with PCA implementation in the *scikit-learn* package [59].

Reactions classified as “Amino acid metabolism” in the inset of Figure 1b are collected from the following model subsystems: “Alanine and Aspartate Metabolism”, “Arginine and Proline Metabolism”, “Cysteine Metabolism”, “Glutamate Metabolism”, “Glycine and Serine Metabolism”, “Histidine Metabolism”, “Methionine Metabolism”, “Threonine and Lysine Metabolism”, “Tyrosine, Tryptophan, and Phenylalanine Metabolism”, “Valine, Leucine, and Isoleucine Metabolism”.

### Production analysis of GEMs

Production analysis was used to determine whether endosymbiont GEMs can produce metabolic precursors required to produce new endosymbiont biomass. First, the models were curated applying a “greedy gapfilling” protocol:

1. All transport reactions were removed from the draft model.
2. All reactions from iML1515-tRNA GEM that either do not have a GPR associated (“spontaneous”) or contain a *pseudo*-GPR *s0001* were added to the models.
3. For GEMs of *Blochmanniella* and *Buchnera*, reaction DHQTi (3-dehydroquinate dehydratase) was added. The enzyme essential for aromatic amino acid synthesis and has not been captured by homology search, but EC 4.2.1.10 has been annotated using Prokka [60].
4. For every metabolite that is present in the cytosolic compartment, metabolites in periplasmic- and extracellular compartments were added.
5. Exchange reactions were added for all metabolites in the extracellular compartment.

A list of 42 intermediates from central carbon metabolism, and major biomass components have been compiled (Table S3). The base medium included simple carbon sources (either glucose or a combination of organic acids) together with inorganics (oxygen, ammonium, inorganic phosphate, sulfate). In different scenarios, individual- or sets of amino acids were supplemented to the base medium. Production analysis was performed for every metabolite of interest as follows:

1. A sink reaction was created for the cytosolic metabolite of interest.
2. Flux through the sink reaction was maximized using flux balance analysis (FBA).
3. Lowest absolute flux values through the metabolic network were determined using parsimonious FBA.
4. Metabolite was rendered produced if the absolute flux through sink reaction was higher than the absolute flux through the exchange reaction of that metabolite.

### Manual curation and simulation of GEMs

Reaction and metabolite annotations were taken from either the draft models and/or the BiGG database [61]. For manually curated GEMs, the biomass equation of the template model (iML1515-tRNA) was used. The GEMs, together with custom scripts for reconstruction, semi-automatic- and manual curation of the models, are available on Zenodo (see Data availability). All GEMs were simulated using CBMPy 0.8.2 [62] under Python 3.9 environment with IBM ILOG CPLEX 22.1.1 linear program solver.

## Abbreviations

GEM: genome-scale metabolic model
ORF: open reading frame
str: strain

## Data availability

All scripts, models, and outputs of analyses, and materials for reproducing the Figures of this manuscript, are available on Zenodo: https://doi.org/10.5281/zenodo.15118322 [63].

## CRediT Author Role Statement

Pranas Grigaitis: conceptualization, methodology, software, data curation, formal analysis, investigation, validation, visualization, writing – original draft, writing – review & editing.

Bas Teusink: funding acquisition, supervision, formal analysis, validation, writing – review & editing. Adria C. LeBoeuf: funding acquisition, project administration, supervision, formal analysis, validation, writing – review & editing.

## Acknowledgements

This research was funded by Human Frontier Science Program Organization (Ref. No: RGP0023/2022, https://doi.org/10.52044/HFSP.RGP00232022.pc.gr.153615; to A.C.L and B.T.). We thank SURF (www.surf.nl) for the support in using the Dutch National Supercomputer Snellius, and Blanka J. Izsó for her input on the reconstruction of draft GEMs.

## Notes

### Competing Interest Statement

The authors have declared no competing interest.

