## Supplementary Material for "A stochastic ratchet during genome streamlining can commit insect endosymbionts to a parasitic or mutualistic fate"

### This file contains:

**Supplementary Figures** 2

**Supplementary Tables** 11

### Supplementary Figures

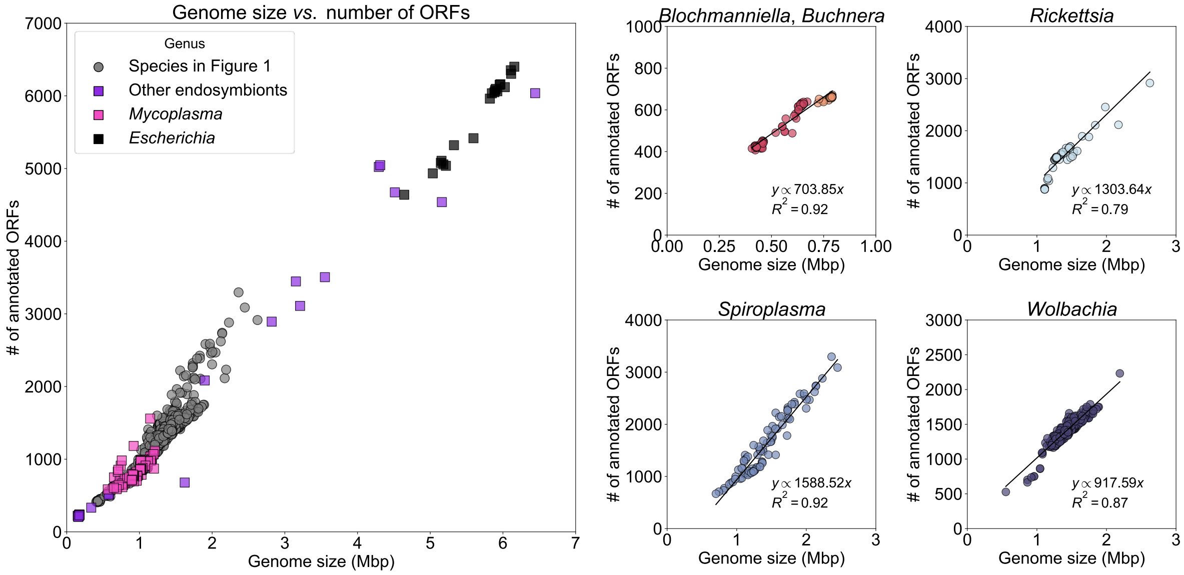


**Figure S1. The number of annotated open reading frames (ORFs) linearly increases as a function of genome size**. **a.** Genome metrics for 548 endosymbiont genomes, including genomes of genus *Sodalis*. Inset: the number of collected genomes per genus. Metrics of 20 genomes of free-living bacterium *Escherichia coli*, and available genomes for multiple endosymbiotic bacteria (except synthetic genomes of *Mycoplasma*) have been included for comparison. **b-e.** Metrics of *Blochmanniella* and *Buchnera* genomes (**b**) or other genera individually (**c-e**). Linear regression fits, slopes and determination coefficient ($R^2$) values were computed using *sklearn* package in Python.

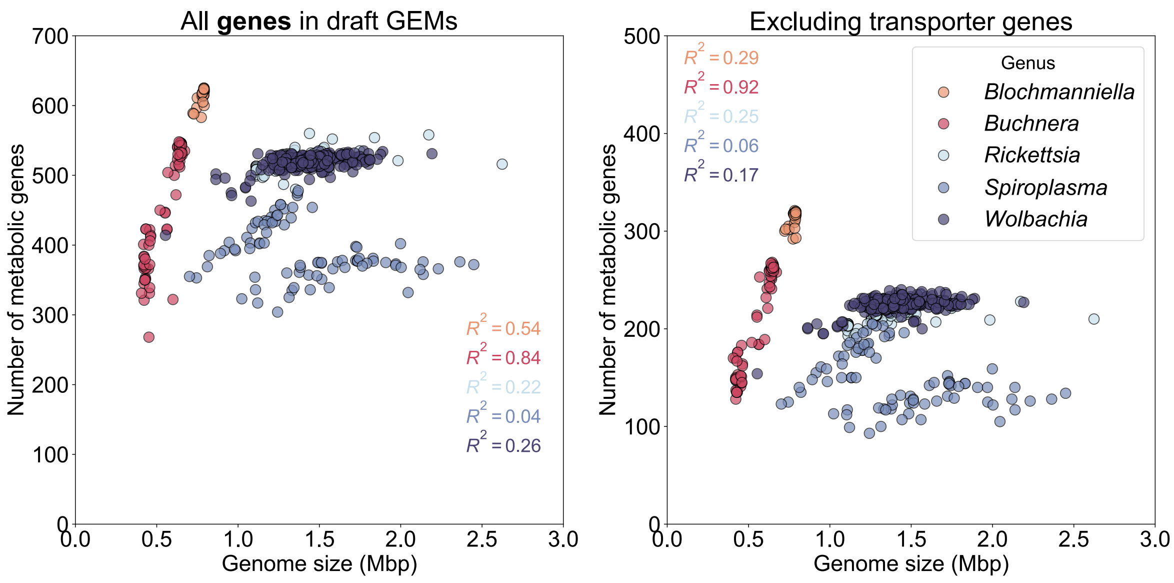


**Figure S2. Correlation of genome size and number of genes in GEMs.**

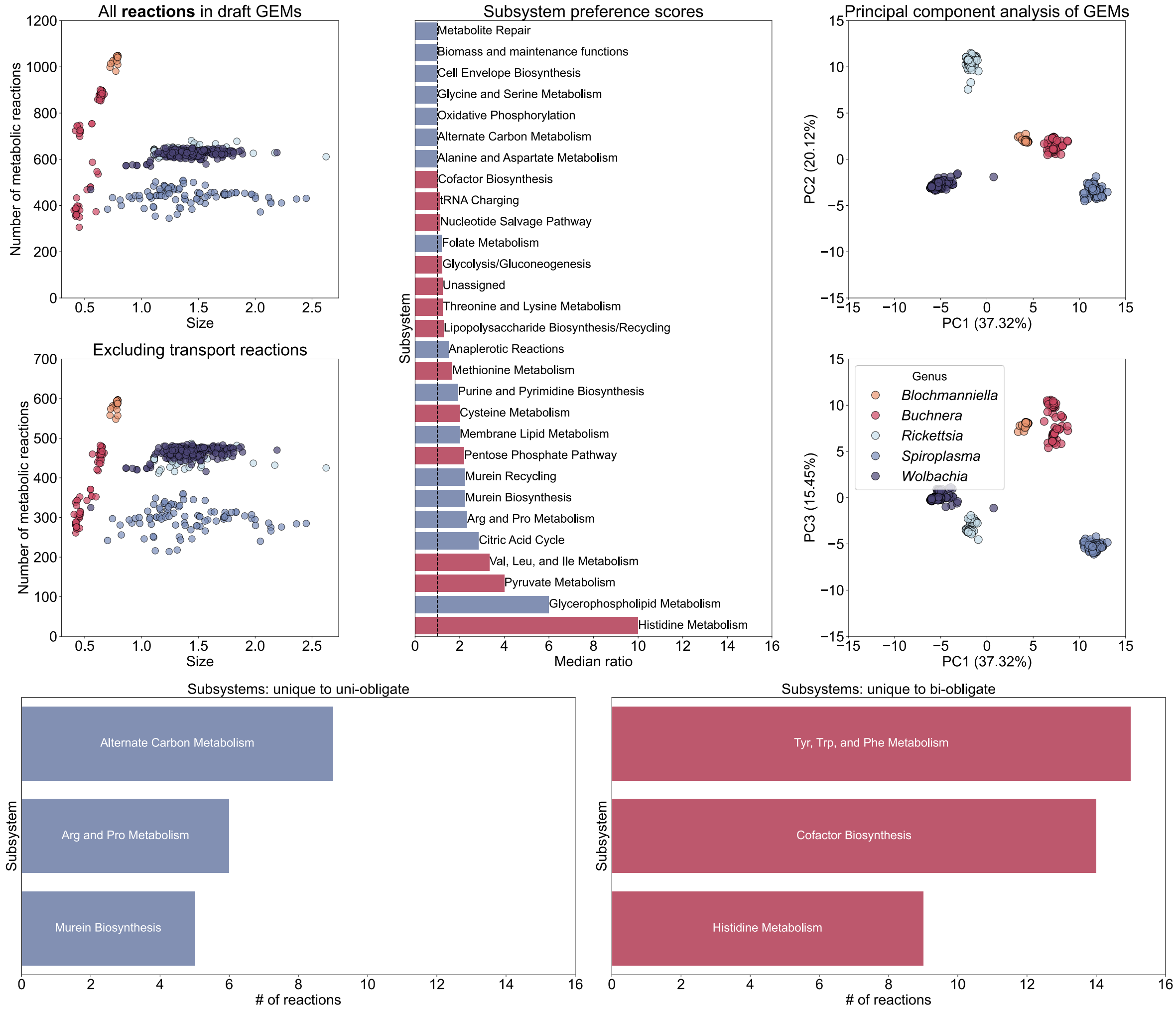


**Figure S3. Comparison of reaction content in GEMs.**

**Top panels:**

**Left:** the number of metabolic reactions in draft GEMs including transport reactions (top) and after correction (bottom). Figure 1b of the Main Text contains the values **after** correction. **Middle**: full output the preference scores for subsystems in GEMs. A median ratio of 1.0 corresponds to no preference between the groups (black dashed line). Blue and red colors indicate subsystems preferred in GEMs of uni- or bi-obligate endosymbionts, respectively. **Right:** principal component analysis of the reactions sets in draft GEMs (see Methods).

**Bottom panels:** Top 3 subsystems containing reactions only in GEMs of uni-obligate (left) or bi-obligate (right) endosymbionts.

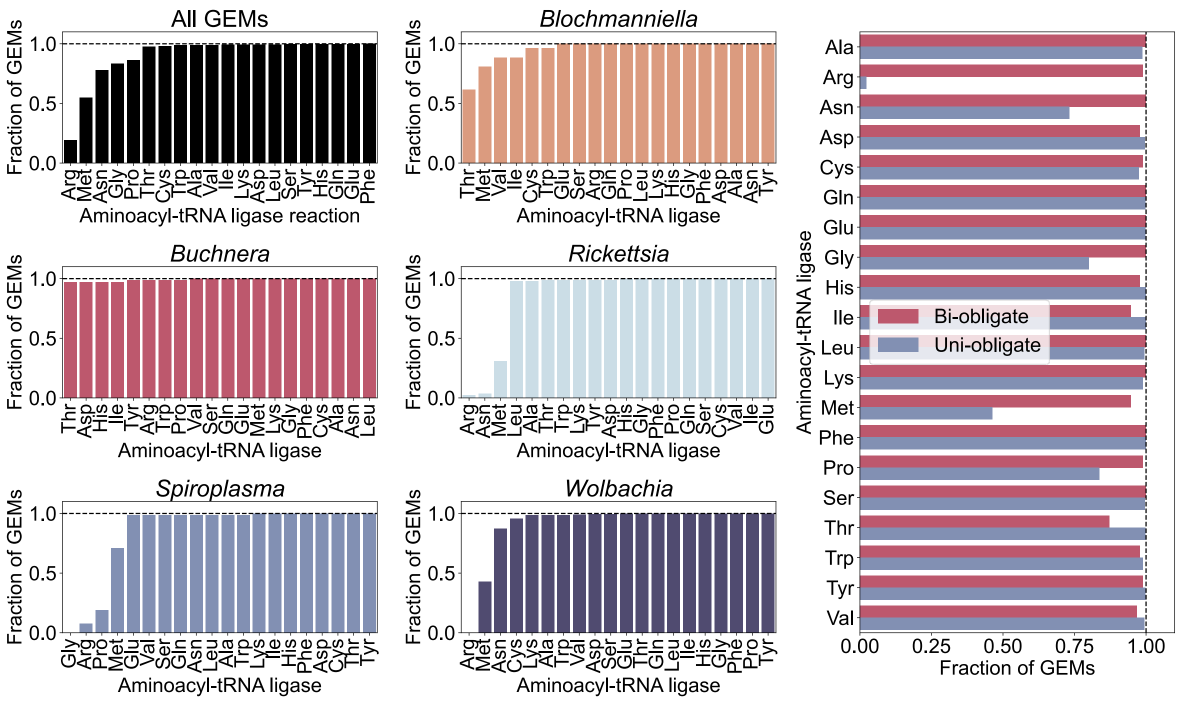


**Figure S4. Methionine-tRNA and arginine-tRNA ligase reactions are missing in many GEMs.** Distribution of aminoacyl-tRNA reactions across all endosymbiont GEMs. Reactions in vertical bar plots are ranked by the fractions of GEMs in the genus containing that reaction.

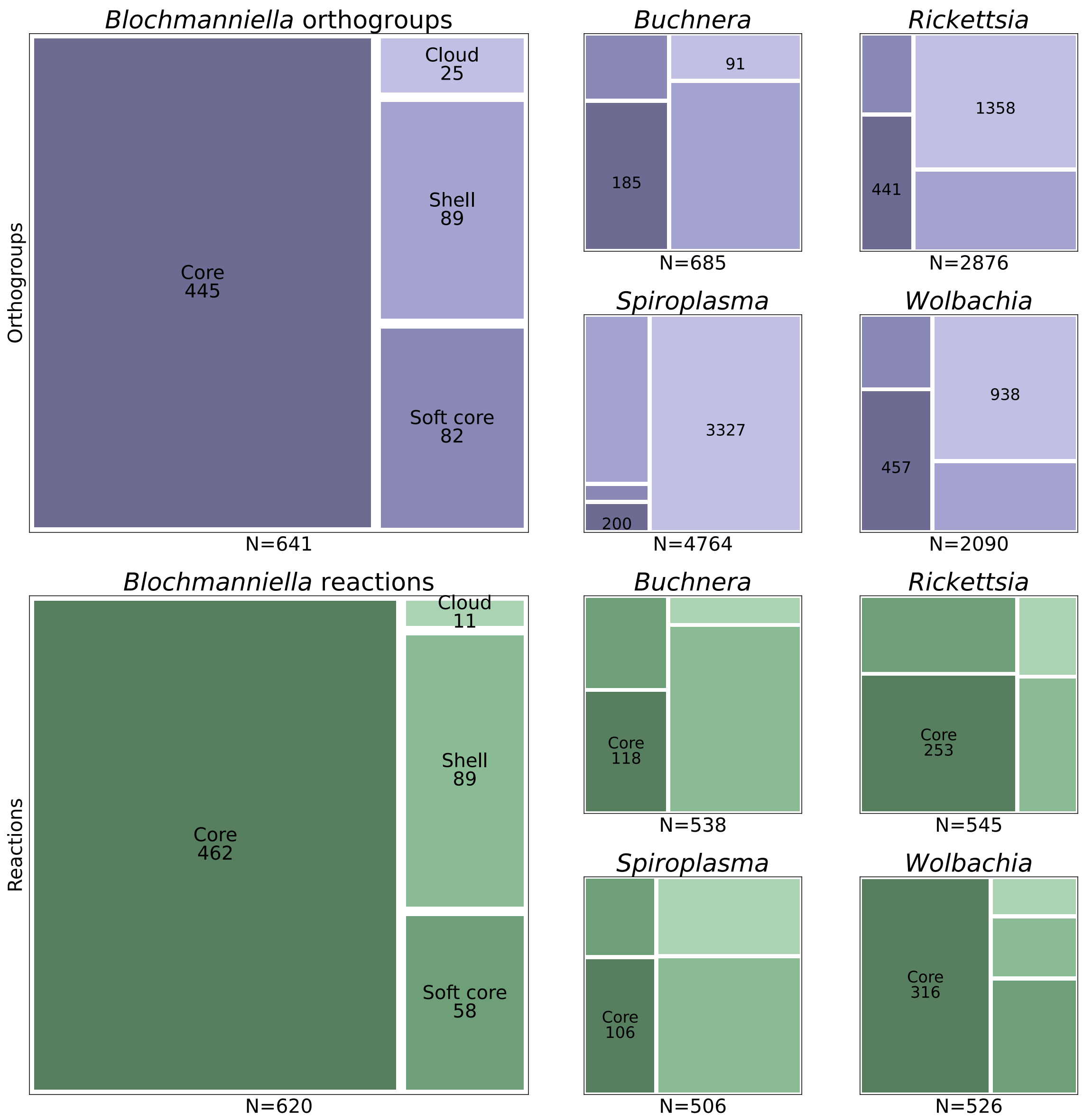


**Figure S5. Endosymbiont genomes and GEMs show high variability within same genus.** Classification of detected gene orthogroups (top) and reactions in pan-GEMs (bottom) according to their occurrence across genomes and GEMs of the genus (see Methods for details). Genes and reactions present in ≥99% genomes/models are marked as core, ≥95% - soft core, and ≥15% - shell. The rest of genes are annotated as cloud. The total number of orthogroups/reactions is listed at the bottom of each treemap.

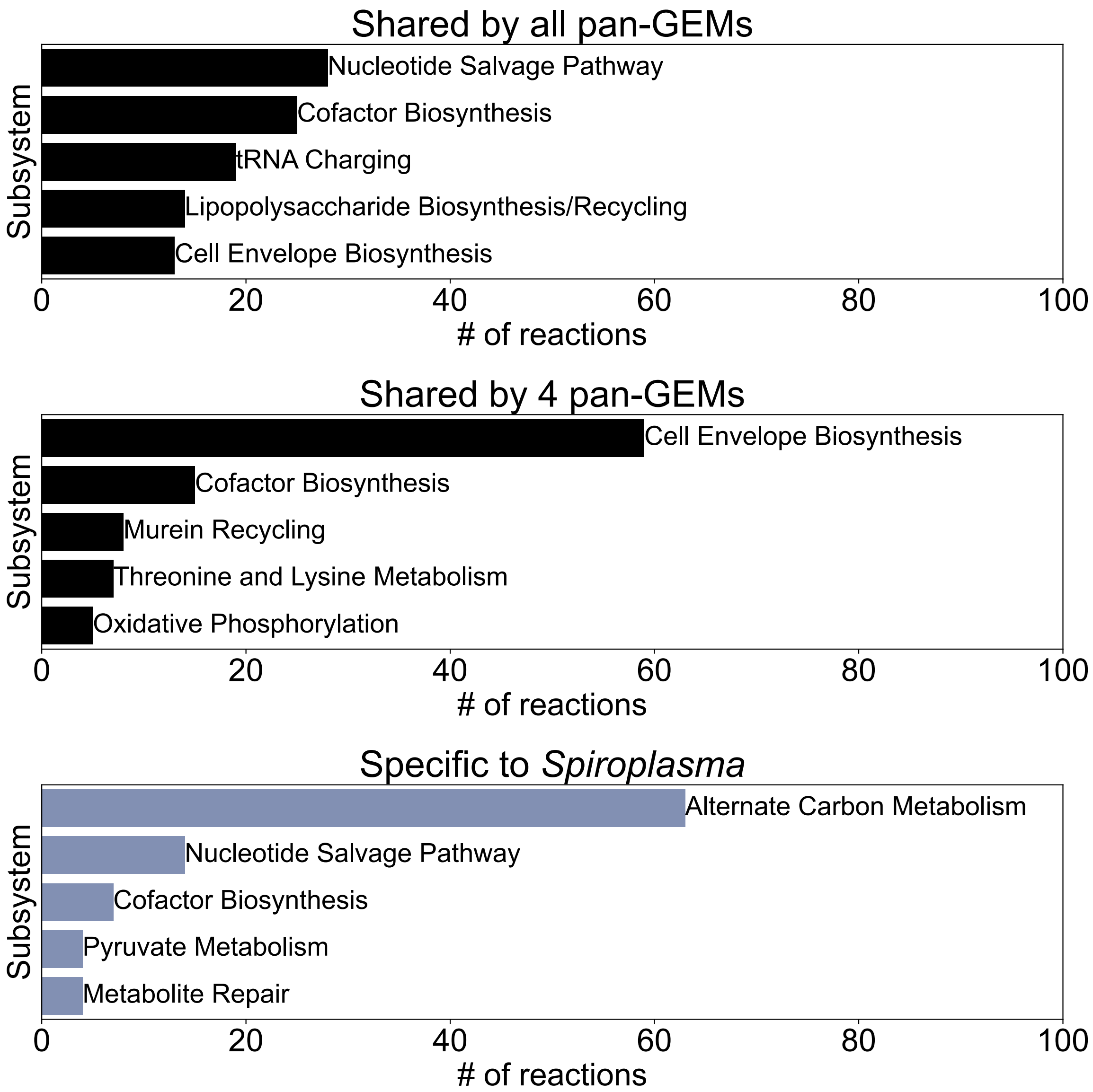


**Figure S6. Endosymbiont pan-GEMs share reactions for resource utilization. a.** Reaction overlaps in pan-GEMs of all 5 endosymbiont pan-GEMs. **b-c.** Metabolic pathways containing most reactions in represented in one of the in pan-GEMs subsets. Some subsystems' names are truncated/altered for clarity.

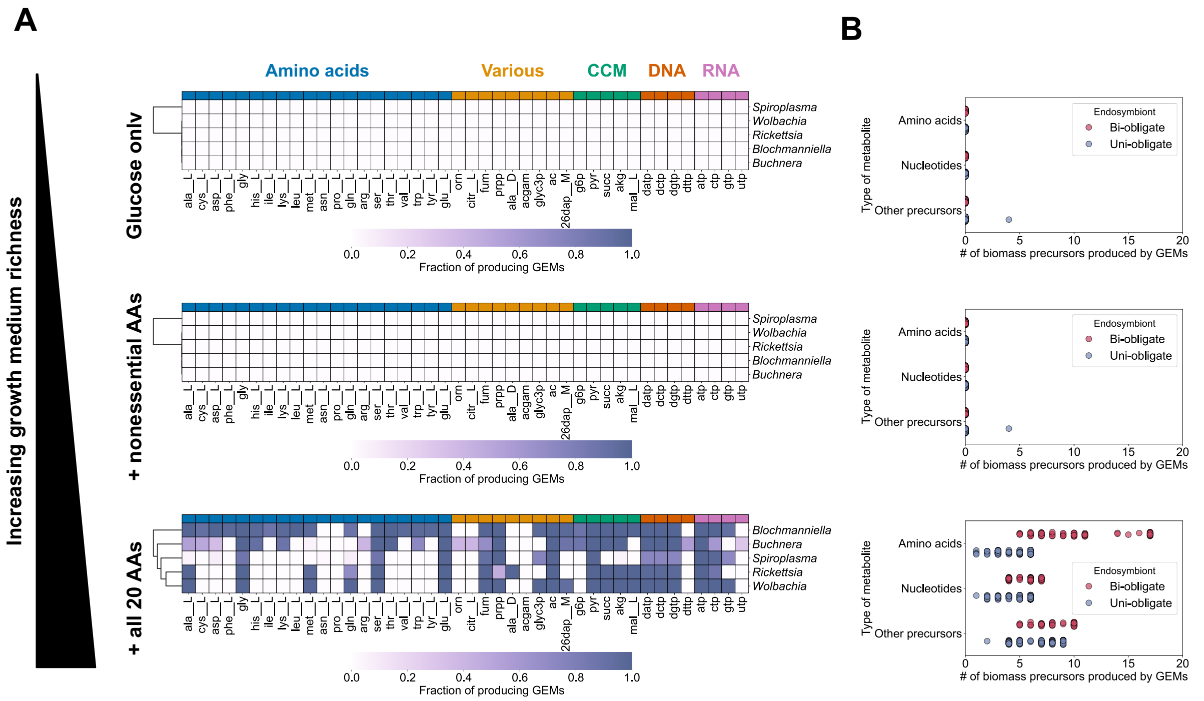


**Figure S7.** Production analysis for endosymbiont GEMs using glucose as the carbon source, instead of set of organic acids as done in Figure 2 of the main text.

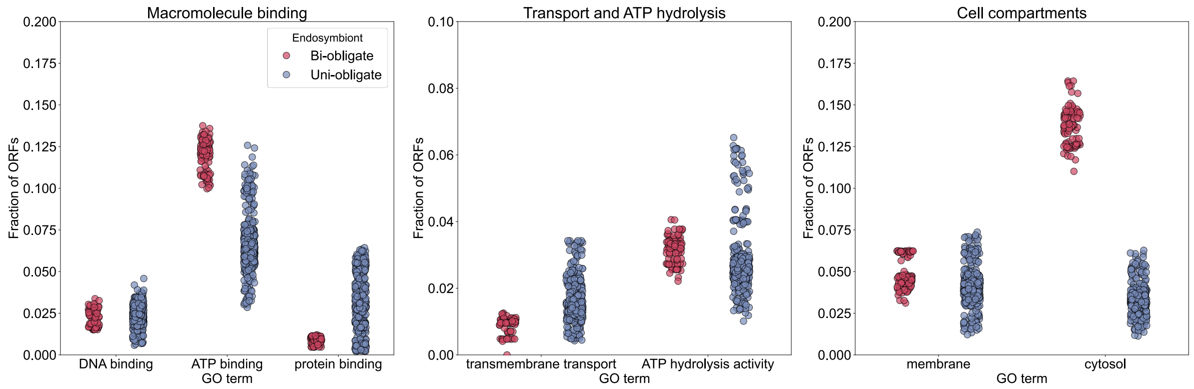


**Figure S8. Distribution of genes in endosymbiont genomes based on their gene ontology (GO) terms normalized to the total number of ORFs.** GO terms related to macromolecule binding (left), cell function and transport processes (middle), and compartmentalization of proteins (right) are the same as in Figure 3 of the Main Text.

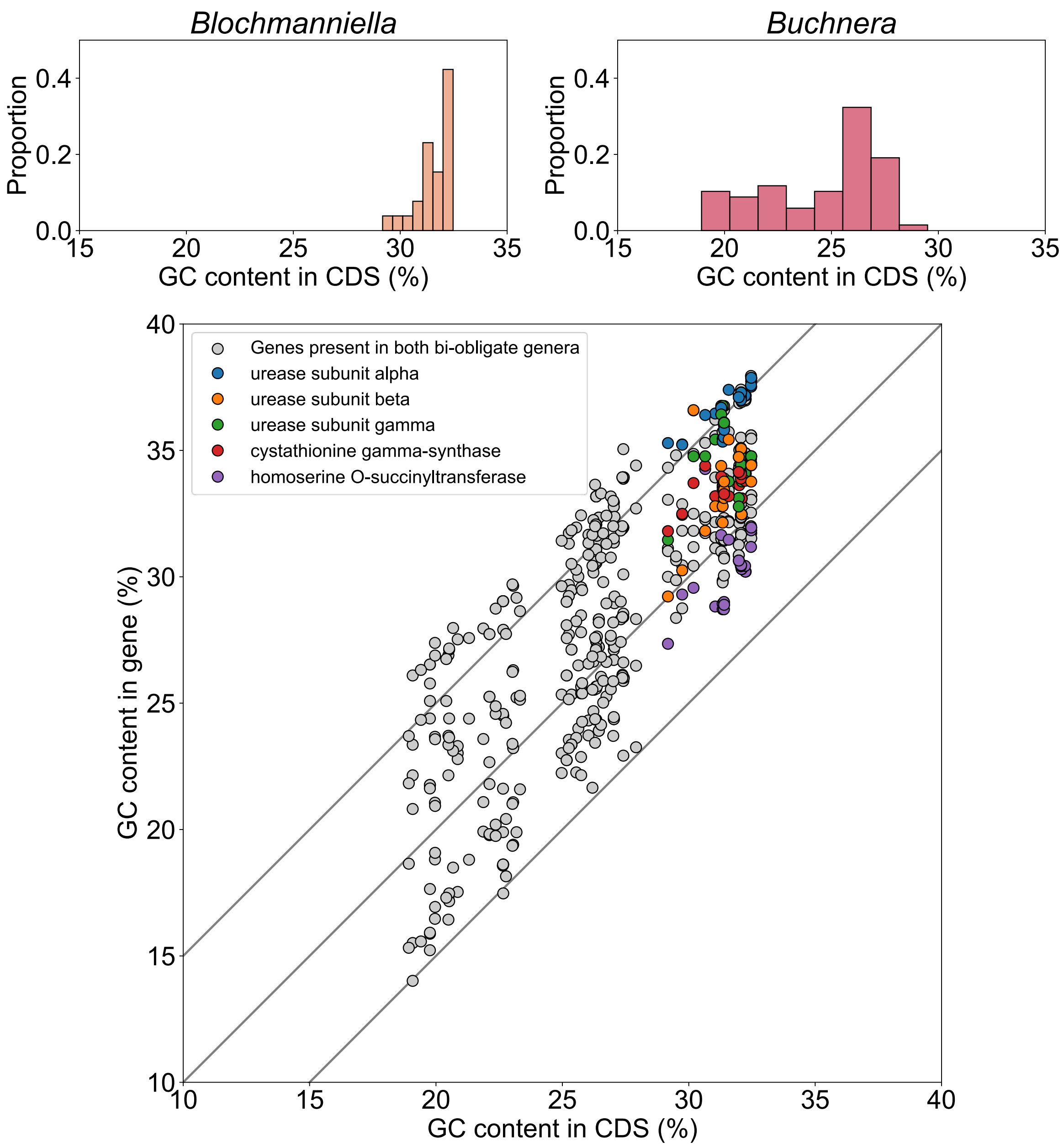


**Figure S9.** The nucleotide composition of genes unique to *Blochmanniella* does not suggest horizontal gene transfer as their origin. **Top:** the GC content in all coding sequences (CDS) of *Blochmaniella* and *Buchnera*. **Bottom:** the relationship between the GC content in all CDS vs. the CDS of representative genes encoded only by *Blochmaniella* genomes. Auxiliary lines represent ±5% deviation from the GC content in CDS.

### Supplementary Tables

**Table S1.** Statistics of genomes collected for different endosymbiont genera.

| Genus | N | Median genome size (Mbp) | Median no. of ORFs |
|---|---|---|---|
| *Blochmanniella* | 26 | 0.786 | 661 |
| *Buchnera* | 68 | 0.584 | 540 |
| *Rickettsia* | 88 | 1.280 | 1476 |
| *Sodalis* | 6 | 4.412 | 4847 |
| *Spiroplasma* | 90 | 1.354 | 1416 |
| *Wolbachia* | 270 | 1.394 | 1392 |

**Table S2.** Comparison of omnipresent orthogroups vs. the number of annotated ORFs in the smallest genomes per endosymbiont genera.

| Genus | Omnipresent orthogroups | Minimal # of ORFs |
|---|---|---|
| *Blochmanniella* | 445 | 663 |
| *Buchnera* | 185 | 407 |
| *Rickettsia* | 441 | 876 |
| *Spiroplasma* | 200 | 668 |
| *Wolbachia* | 252 | 527 |

**Table S3.** Selected metabolites for production analysis.

| Category | Metabolites |
|---|---|
| Amino acids | All 20 proteinogenic amino acids |
| Biosynthetic precursors | L-ornithine, L-citrulline, fumarate, D-alanine, phosphoribosyl pyrophosphate, N-acetyl-D-glucosamine, meso-2,6-diaminoheptanedioate, glycerol 3-phosphate, glucose 6-phosphate, pyruvate, succinate, alpha-ketoglutarate, L-malate |
| Ribonucleotides | ATP, CTP, GTP, UTP |
| Deoxyribonucleotides | dATP, dCTP, dGTP, dTTP |

**Table S4.** Metrics of draft- and manually curated endosymbiont GEMs. Asterisk denotes an assembly suspended by RefSeq and thus not included in the previous analyses. n/a: not available.

| Species | RefSeq assembly ID | Draft model | | Curated model |
|---|---|---|---|---|
| | | Reactions | Genes | Reactions |
| *Ca B. floridana* | GCF_000043285.1* | 1028 | 318 | 1091 |
| *B. aphidicola* | GCF_001700895.1 | 874 | 264 | 942 |
| *R. conorii* | GCF_000007025.1 | 635 | 242 | n/a |
| *S. kunkelii* | GCF_001274875.1 | 417 | 139 | n/a |
| *Wolbachia sp.* | GCF_016584425.1 | 636 | 255 | n/a |
